# A Self-Supervised Learning Approach for High Throughput and High Content Cell Segmentation

**DOI:** 10.1101/2024.05.29.596446

**Authors:** Van Lam, Jeff M. Byers, Michael Robitaille, Logan Kaler, Joseph A. Christodoulides, Marc P. Raphael

## Abstract

In principle, AI-based algorithms should enable rapid and accurate cell segmentation in high-throughput settings. However, reliance on large datasets, human input, and computational expertise, along with issues of limited generalizability and the necessity for specialized training are notable drawbacks of nominally “automated” segmentation tools. To overcome this roadblock, we introduce an innovative, user-friendly self-supervised learning method (SSL) for pixel classification that requires no dataset-specific modifications or curated labelled data sets, thus providing a more streamlined cell segmentation approach for high-throughput and high-content research. We demonstrate that our algorithm meets the criteria of being fully automated with versatility across various magnifications, optical modalities and cell types. Moreover, our SSL algorithm is capable of identifying complex cellular structures and organelles which are otherwise easily missed, thereby broadening the machine learning applications to high-content imaging. Our SSL technique displayed consistent F1 scores across segmented images, with scores ranging from 0.831 to 0.876, outperforming the popular Cellpose algorithm, which showed greater variance in F1 scores from 0.645 to 0.8815, mainly due to errors in segmentation. On average, our SSL method achieved an F1 score of 0.852 ±0.017, exceeding Cellpose’s average of 0.804 ±0.08. This novel SSL method not only advances segmentation accuracy but also minimizes the need for extensive computational expertise and data security concerns, making it easier for biological researchers to incorporate automated segmentation into their studies.

## Introduction

Today, 384 and 1536 well plate layouts are quickly replacing the past norm of 96 well plates and becoming commonplace in high-throughput assay applications such as drug impact and toxicity assessment, expanding the range of cell lines, treatments, phenotypes and surfaces for a given experimental dataset ^1–3^. To keep pace, the application of machine vision for cell segmentation has grown remarkably to avoid time consuming and intensive manual equivalents. Automated cell segmentation is seen as a cornerstone for high-throughput studies, highlighting the need for software which can facilitate the accurate assessment of cellular morphology. The applications are broad for automated cell segmentation due to the need for swift and precise analysis in high-throughput contexts for drug development, real-time gene-expression tracking, and cell therapy ^4–7^ to name a few. Despite numerous reports of success with current automated segmentation tools, the application of deep learning in cell segmentation still encounters challenges, including a significant demand for data, the need for human intervention and biases within, limited generalizability and specialized training.

Deep convolutional neural networks (CNNs) have gained prominence in deep learning-based cell segmentation, showcasing the efficiency and accuracy of UNET ^8–12^ and Mask R-CNN ^13^ in cell and nucleus segmentation. This technique interprets segmentation as a task of identifying cell boundary pixels, compelling the model to produce a ‘distance map’ for each pixel to estimate its likelihood of being at a cell border. In high-throughput assays, a diverse dataset capturing a broad spectrum of cell morphologies is crucial for CNNs to accurately segment individual cells. However, the extensive requirement for data training and annotation restricts the application of CNNs in high-throughput studies with limited replication and presents a significant hurdle for researchers who are unable to expand their studies without additional training data. A recent study highlights that effective CNN model training requires a substantial, high-quality pre-training dataset consisting of 1.6 million cells, presenting the challenge of applying CNN methods in segmenting specialized high-throughput studies due to the sheer scale of data needed ^14^. Other techniques like pixel classification are preferred for images with low cell density when it involves labeling each pixel in an image as a distinct class (e.g., cell versus background)^15–18^. High fidelity segmentation is essential to further downstream tasks such as declumping and organelle identification using post-processing algorithms like watershed ^19^, active contour ^20^ or threshold-based segmentation^21^.

According to the 2018 Data Science Bowl, the top three performance CNN models, namely U-Nets, fully connected FPNs, and Mask R-CNN demand high computational expertise for segmentation tasks, resulting in a significant obstacle to biologists who are the primary end-users of these models ^9^. Furthermore, it is of paramount importance to expand the training datasets and enhance their ability to generalize across different studies that go beyond competition scenarios. Given that scientists frequently resist expanding extensive imagery datasets for additional training, an innovative strategy in bio-imaging platforms adopted by CNNs such as Cellpose 2.0 ^22^ and DeepCell ^23^, involves incorporating a “human-in-the-loop” feature to aid in segmentation. This feature enables users to adjust crucial cell parameters manually or undertake the segmentation of specific cells of interest for a more targeted training set. Nevertheless, this approach introduces additional labor and time, diverging from the goal of completely automated high throughput imaging. Furthermore, users may have to offer numerous manual segmentation references for each condition in the high-throughput assay, potentially increasing bias. Without objective and reliable cell-from-background segmentation, critical downstream tasks such as declumping and morphological measurements are negatively impacted.

In addition to the increasing focus on 384 and 1536 well plates, high throughput imagers are now equipped with high magnification, immersion objectives for high-resolution imaging (*e.g*. objectives with 50x-60x magnification and high numerical apertures). This synergy of high-throughput and high-resolution imaging can significantly enhance the comprehensive understanding of cell phenotype, covering aspects such as sub-cellular structure ^23,24^, molecular expression ^25^, and organelle distribution ^26^. For instance, high-resolution imaging’s potential enables scientists to quantify cell membrane convexity and concavity, offering insights into the interactions between cytoskeleton and substrate environment ^27^. Despite these advances, current studies often overlook the meticulous analysis of fine cell structure due to the limitations of study size and a lack of annotated training data required for high content automated segmentation. Consequently, biomedical researchers are frequently compelled to choose between image quality and sample size. The future of cell imaging must bridge the gap between high-resolution and high-throughput methodologies by advancing towards their integration, making fully automated segmentation an essential analytical tool. With this capability, researchers can unlock a far more comprehensive understanding of cellular behavior captured from detailed structural nuances of vast datasets, generated from imaging thousands to millions of cells across various conditions, materials, and time points.

To address these challenges, we present a novel self-supervised learning (SSL) methodology for pixel classification tasks, aimed at automating single cell segmentation for both high-content and high-throughput data studies. This user-friendly and robust approach is completely automated, eliminating the need for dataset-specific adjustments and data annotation in the training process. Our previous segmenting method utilized optical flow vector fields generated by consecutive images from live cell time series data sets. Building on this, we demonstrate a method *applicable to both live or fixed cell imagery*, achieved by augmenting a single image and then employing self-supervised learning techniques to both the original and augmented images. Specifically, we employ a Gaussian filter to create an augmented image from an original input image, then calculate optical flow from the augmented image. Optical flow vectors are used as a basis for self-labelling pixel classes (‘cell’ vs ‘background’) to train a classifier. We demonstrate that this SSL approach achieves complete automation while also capturing high content information with a robustness that extends across a range of magnifications and optical modalities. As a result, we present this algorithm as an enabling technology that applies across disciplines, from exploratory cell biology to targeted biomedical research studies.

## Materials and Methods

### Cell culture and microscopy

All mammalian cells were grown in DMEM (ATCC, #30-2002) enriched with 10% fetal bovine serum (ATCC, #30-2020) at 37°C in a 5% CO_2_ atmosphere. Cell treatments and growth conditions included a range of environments including off-the-shelf glass-bottomed dish culturing, or fibronectin treated surfaces. Detailed culture methodologies and immunofluorescence staining protocols are described in the references ^17,18,28^. Microscopy information for each cell type, including the mode of microscopy, magnification, numerical aperture, and camera type is provided in Supplementary Note 1.

### Self-Supervised Learning

The foundational principles of employing optical flow for SSL-based live cell image analysis were elaborated in earlier publications in which two images from consecutive time points were utilized for labeling and training ^17,18^. Here we broaden the approach to single or fixed-cell imagery, requiring only a single input image for the entire segmentation process. This is accomplished by applying a Gaussian filter (σ=0.1) to the original image to generate an augmented image, which serves as an input for calculating a Farneback optical flow displacement field. The Farneback algorithm determines optical flow of a moving object by calculating the displacement field at multiple levels of resolution, starting at the lowest and continue until convergence. The Gaussian-filtered augmented image is synergistic to this approach in that it is less noisy while retaining the optical flow requirement of preserving overall image intensity. Less noise aids the Farneback algorithm in more effectively utilizing lower-resolution images to accurately segment objects.

Cells typically have higher entropy (or texture) than the background due to complex internal structures such as organelles and the cytoskeleton. We take advantage of this fact by using this entropy difference as a ruler for self-tuning and optimizing the optical flow thresholds that label pixels as either ‘cell’ or ‘background’. This self-tuning phase is termed unsupervised learning as it operates without a predefined training library. Following the unsupervised learning stage, masks for cell and background are created, each being multiplied by the original image. The resultant image products are analyzed to extract features such as entropy, gradient, and intensity from every labelled pixel.

The process transitions to supervised learning when feature vectors from each pixel, along with their corresponding labels (‘cell’ and ‘background’), are aggregated and introduced to a Naïve Bayes Classifier, employing a 10-fold split for validation. Subsequently, a model is derived and applied to the original image, culminating in the final segmentation of cells. This methodology underscores a comprehensive approach, integrating both unsupervised and supervised learning phases to achieve precise cell segmentation without the necessity for an extensive training dataset or manual parameter tuning. It is intrinsically adaptive in that a new model is created for each image.

### Manual segmentation for F1 score evaluation

An in-house MATLAB script was developed to facilitate manual segmentation, utilizing the drawpolygon function. The resulting cell masks were then saved and employed to compute the F1 score for both the Cellpose and SSL techniques. The F1 score, a harmonic mean of precision and recall, is defined as

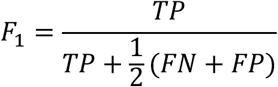

Where TP, FN and FP are abbreviations for true positive, false negative and false positive, respectively.

This approach enables a quantitative comparison of the segmentation accuracy between the Cellpose and SSL methods, providing a measure of their performance in terms of both precision (the proportion of true positive results in all positive predictions) and recall (the proportion of true positive results in all actual positives).

## Results

### Self-supervised learning approach for segmenting diverse microscopy modes across multiple cell types

We first aim to highlight the compatibility of our SSL code with high throughput imaging in that robust cell segmentation takes place from single images in the absence of both curated training libraries and manual parameter settings. Our SSL method demonstrated versatility by working at different resolutions and with different image modalities spanning from label free to fluorescent modalities (Figure 2 a-e). Representative images segmented by SSL varied from mammalian cell types to fungi, processed in an automated fashion without the need for a curated training library or manual filters. Label-free examples includes breast cancer cells MDA-MB-231 imaged by phase contrast at 10x (Figure 2a) and bright-field at 40x (Figure 2b); differential interference contrast (DIC) images of MDA-MB-231 at 63x (Figure 2c.i) and *S. cerevisiae* (Figure 2c.ii); human fibroblast Hs27 imaged by interference reflection microscopy (IRM) at 40x (Figure 2d); epifluorescence images of GFP-labeled A549 cells (previously analyzed as live cell imagery in references at 40x (Figure 2e.i and ii) at 40x ^17,29^, F-actin and Vincullin-labeled MDA-MB-231 cells (Figure 2e.iii and iv) at 63x. The combined segmentation of both structures F-actin (in red) and Vinculin (in green) appears yellow, allowing for further co-localization analysis of these structures. An additional declumping step was employed in Figure 2c.ii after segmentation by SSL (Supplementary Note 2).

**Figure 1:**
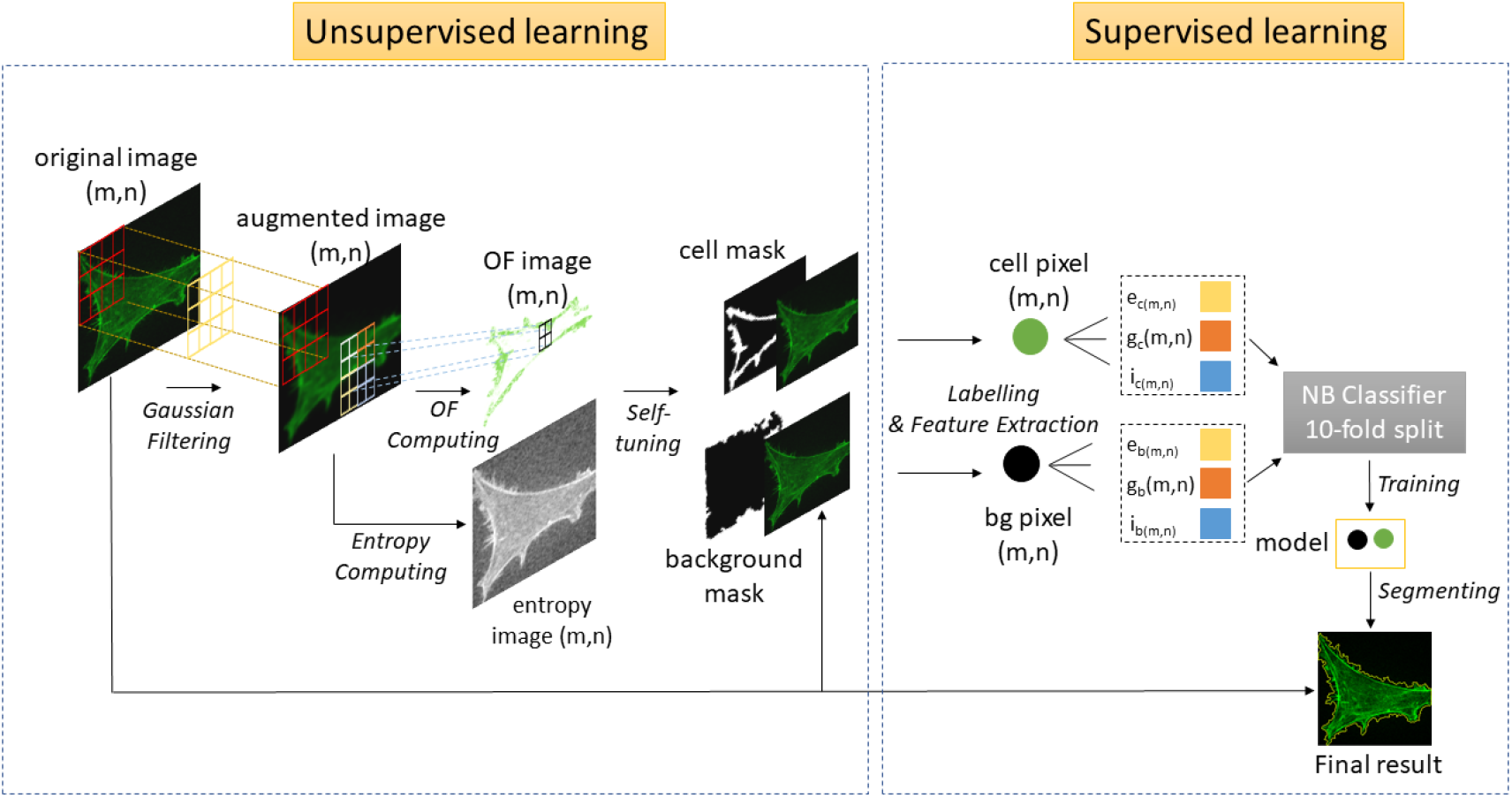
Flow chart of self-supervised learning segmentation. The process starts with generating a augmented image, then applying unsupervised learning via optical flow (OF) on the augmented image to label cell and background (bg) pixels. Supervised learning is performed by extracting labelled (cell or background) pixel features from the augmented image including entropy, gradient and intensity. A Naïve Bayes Classifier with 10-fold split is used for training. The classification model is exported and applied to the original image. This method is repeated with each image ensuring an adaptive process in which every image has a uniquely associated classification model.

**Figure 2:**
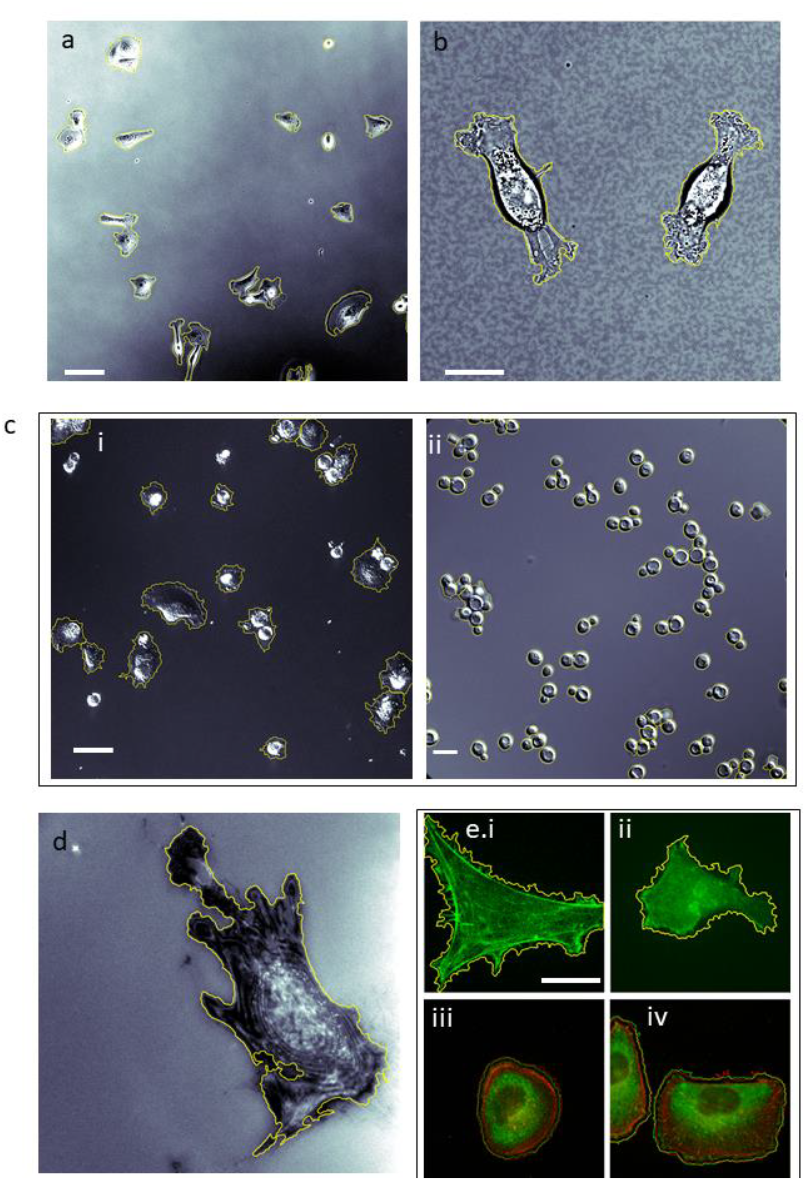
Self-supervised learning segmentation approach for a range of cell types, optical modalities and magnifications. Phase contrast image of (**a**) MDA-MB-231 (10X objective), bright-field image of (**b**) MDA-MB-231 (40X objective), (**c**) DIC image of MDA-MB-231 (i) (20X objective) and *S. cerevisiae* (ii) (63X objective), (**d**) IRM image of Hs27 (40X objective), (**e**) epifluorescence images of (i) A549 cells with lifeAct (GFP-actin conjugate, 100X objective) and (ii) MDA-MB-231 (F-actin and vinculin expression, 63X objective). Yellow border show SSL segmentation result. Scale bar **a**: 50μm, **b**: 10μm, **c**.**i**: 25μm, **c**.**ii**: 5μm, d: 20μm and **e**: 20μm.

### Self-supervised learning for segmenting diverse fluorescent channels across multiple cell treatments

Fluorescent Hs27 cell segmentation was performed on epi-fluorescent and confocal imagery with the cells cultured on different surfaces and stained with DAPI, phalloidin, and anti-Vinculin antibodies (Figure 3). In a completely automate fashion, SSL achieved robust cellular segmentations across different stains: DAPI (Figures 3a.i,iv), phalloidin (Figures 3a.ii, b.i) and vinculin-antibody (Figures 3a.iii, b.ii)); different modalities (epifluorescence vs confocal microscopy); and different resolutions (20x vs 63x). The nuclei were segmented with robust, clear borders, though in some instances, double nuclei were observed which could be explained by cell division (Figure 3a.i). Most individual cell were successfully segmented by SSL via F-actin and Vinculin structures across different imaging setups, demonstrating the segmentation’s adaptability and precision at various resolutions without the need for curated pre-training data sets. In addition, membrane morphologies unique to a given stain could be distinguished upon merging these segmentations (Fig 3b.iii).

**Figure 3:**
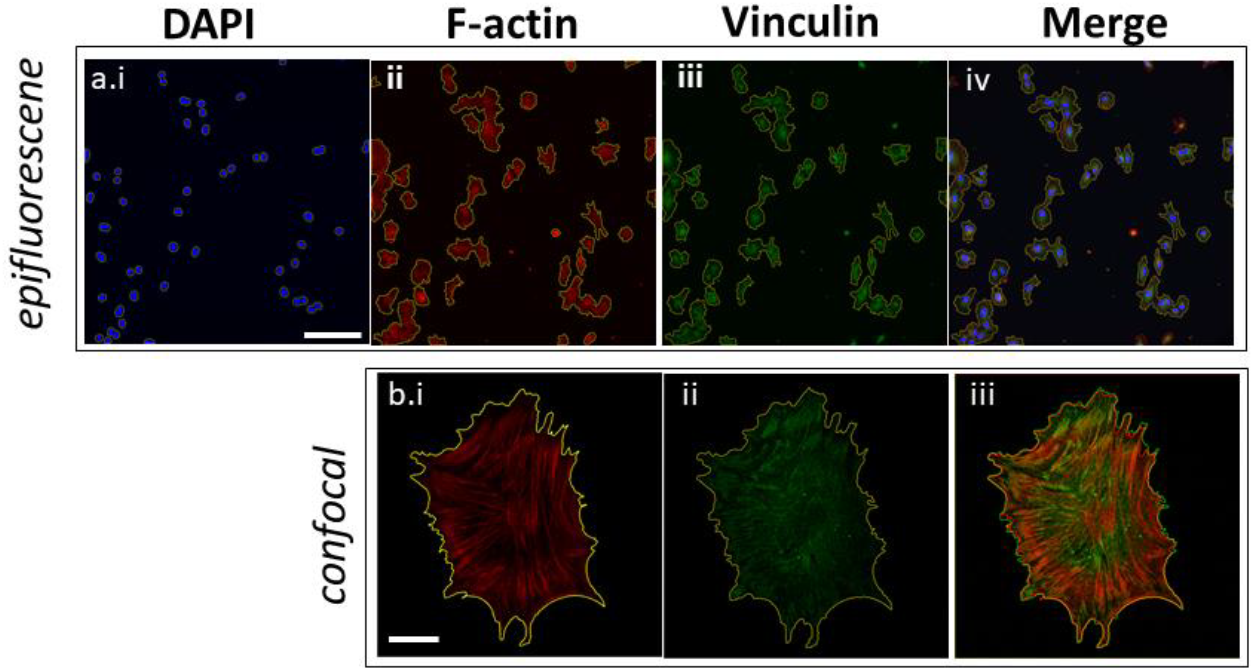
Self-supervised learning segmentation across different cell culture methods, resolutions and cellular structures. Fluorescent images of Hs27 stained for nucleus (DAPI), cytoskeleton (F-actin) and focal adhesions (Vinculin) cultured on (**a**) glass-bottomed petri dishes using a 20x objective (epifluorescence) and (**b**) 63x confocal images of Hs27 cultured on glass-bottomed petri dish stained for F-actin and Vinculin. Scale bar **a**: 50μm, **b**: 10μm.

### Self-supervised learning segmentation approach for quantifying fibronectin’s phenotypic stimuli

Given that most high-throughput experiments are performed on fluorescent data across a variety of treatment conditions, we conducted an analogous experiment by compiling images of fibroblast Hs27 plated atop surfaces treated with different fibronectin concentrations in a single executable run. These surfaces were drop coated with fibronectin of 5, 10, 25, 50, 100 μg/ml, fixed and stained with Vinculin, then imaged using 40x confocal microscopy. Images of Hs27 cells cultured on surfaces treated at different fibronectin concentrations were loaded and processed with SSL in one shot, ensuring the initial self-tuning values for background and cell pixel labelling during unsupervised learning remained consistent. As in previous examples, the need for a training dataset or tuning parameters was eliminated, enabling wide ranging phenotype segmentation in an automated fashion. The entire segmentation process for this particular experiment was completed in 165 seconds, efficiently processing 15 images throughout five conditions. In parallel, the same images were analyzed using the Cellpose CYTO model, with manual input for the object diameter set to 150 pixels that was effective in most cases. Figure 4 presents a direct comparison between the segmentation results of Cellpose (in red) and SSL (in yellow). Our SSL approach consistently delivered robust segmentation outcomes for Hs27 cells, requiring no manual intervention, even with the variations in cell shapes induced by fibronectin treatments. Conversely, the Cellpose model sometimes failed to identify the elongated trailing edges of fibroblasts (Figure 4c, e, h, j) or missed entire cells (Figure 4i). Hs27 morphology varies dramatically between low and high fibronectin concentration and therefore requires an adaptable segmentation approach which our SSL approach achieves by ensuring that every image has a uniquely associated classification model. Indeed, SSL had no difficulty segmenting the completely unique phenotypes associated with low fibronectin treatments of 5 or 10μg/ml (Figure 4a, f) alongside higher concentrations.

**Figure 4:**
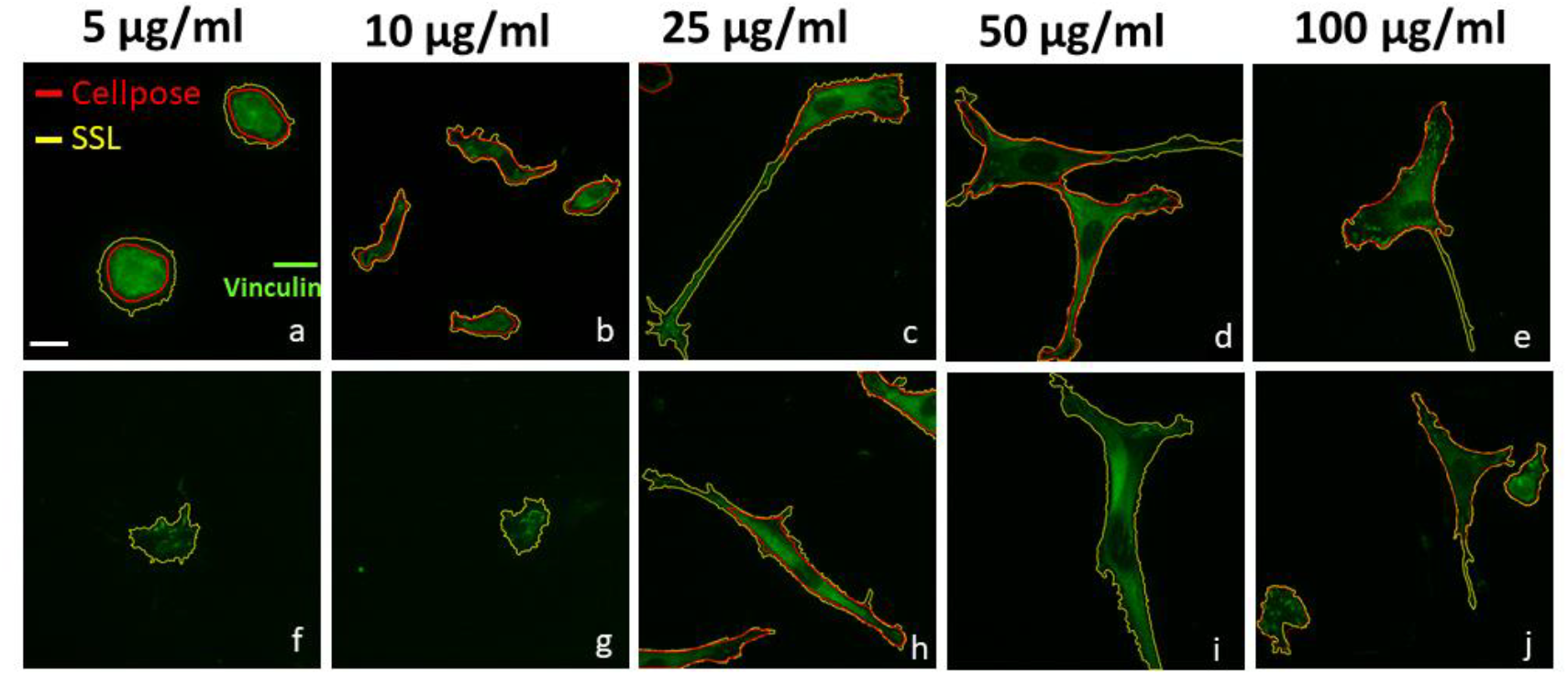
Self-supervised learning segmentation approach in studying the effects of fibronectin concentration on Hs27. Fluorescent images of Hs27 stained for focal adhesion (Vinculin) cultured on glass-bottomed petri dishes using 40x objective. SSL results are outlined in yellow and Cellpose in red. Scale bar **a-j**: 20μm.

### Quantitative analysis of SSL and Cellpose performance on fluorescent segmentation

To better compare the performance of SSL and Cellpose in fluorescent imagery segmentation, another fluorescent datasets of Hs27 cells stained for F-actin, imaged by 10x epi-fluorescence and 60x confocal microscopy, were segmented using SSL, retrained Cellpose (CP) and Cellpose assisted with human-in-loop feature (CP-HiL). Since the data set is relatively small (10 images), we took 7 images for training the new Cellpose model and 3 images for testing the model. The initial segmentation of CP had some inaccuracies on the 10x images (Figures 5 a.i-iii), highlighted by dotted boxes, prompting us to integrate a CP-HiL approach for manual corrections of misclassified segments. The CP-HiL was utilized by combining our in-house 10x F-actin fibroblasts trained by the built-in Cellpose model with manual segmentation to improve performance. Nevertheless, the CP-HiL results were also varied and not improved (Figure 5 b.i-iii). Though Figure 5b.i showed that CP-HiL successfully caught missing cells from the CP method shown in Figure 5a.i, CP-HiL failed to identify other cells or misinterpreted a single cell as two separate entities (Figure 5b.ii,iii). Segmentation with SSL did not overlook any cells or falsely split single cells, though additional declumping algorithms will need to be added as a separate step in future work (Figure 5c.iii).

**Figure 5:**
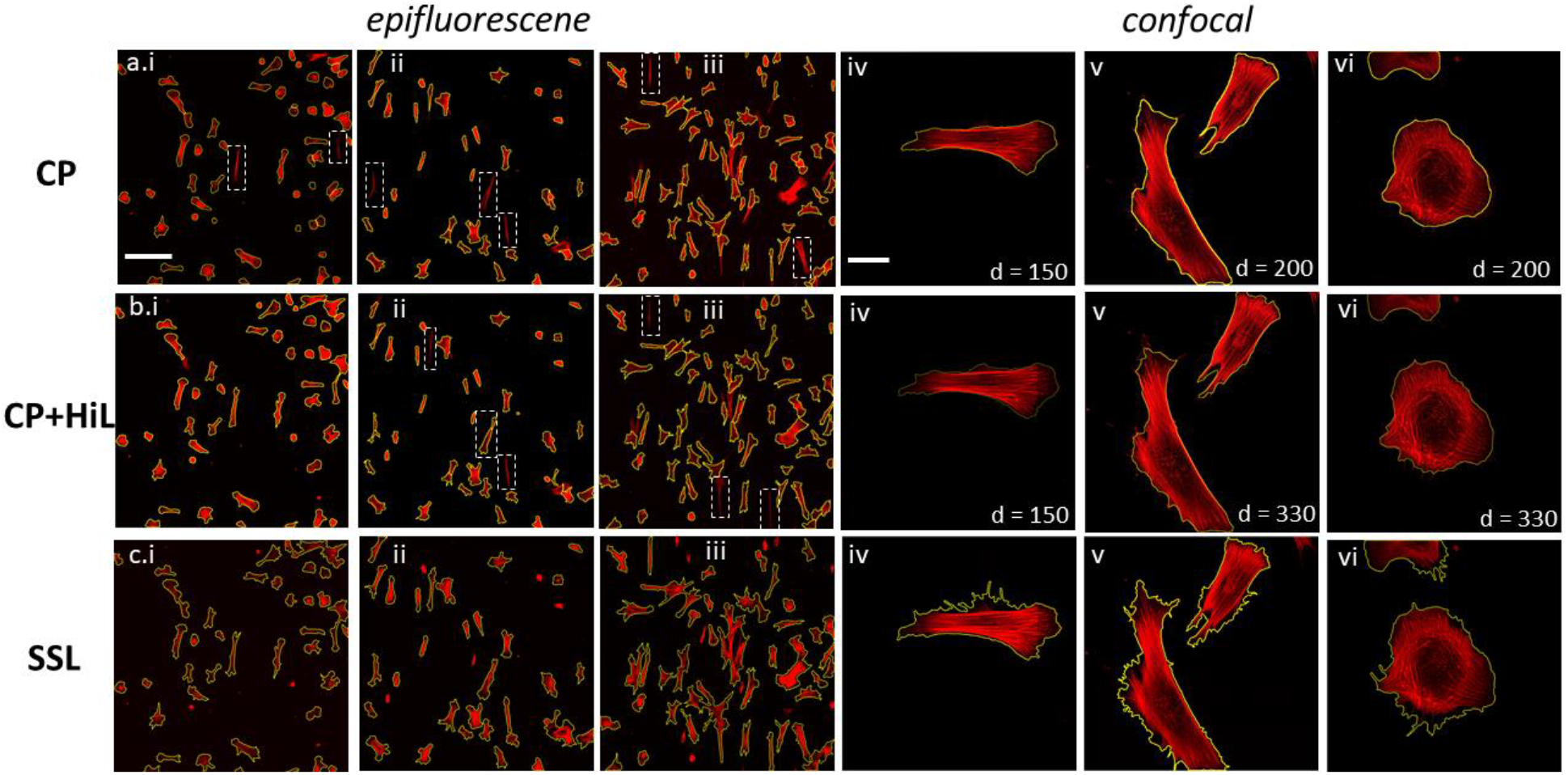
Representative segmentation results on images of Hs27 stained with F-actin by (a) Cellpose (CP), (b) Cellpose with human-in-loop feature (CP+HiL) and (c) self-supervised learning (SSL). Dotted-line boxes indicate cells or region of interest that were lost during segmentation. Epifluorescence images were taken with a 10x objective and confocal images were taken at 63x objective. Scale bar **a, b, c** (**i-iii**): 50μm; **a, b, c** (**iv-vi**): 10μm.

Despite the success of CP and CP-HiL in segmenting Hs27 images taken by confocal microscopy, the two methods required manual tuning of the object diameter specific to each image. For instance, a diameter of 150 pixels was selected for Cellpose methods to segment spread out Hs27 cells shown in Figure 5a.iv and 5b.iv. Nonetheless, the two techniques required different object diameters of 200 and 330 pixels for similar cell shapes shown in Figure 5a.v and 5b.v, respectively. The 200 and 330 object diameter, on another hand, was well-applied on segmenting epithelial-like cells (Figure 5a.vi and 5b.vi), resulting in a lack of consistency and clarity in establishing a standardized procedure for segmenting a single data set. Our SSL approach, in fact, maintained a constant image setting inputs across all images, resolutions, and cell shapes (Figure 5 c.i-vi), simplifying the segmentation process and eliminating the need for manual adjustments. Compared to Cellpose performance on high-resolution 60x images, our SSL captured a broader and more detailed view, potentially revealing fluorescent signals that are difficult to detect with the naked eye, as will be explained in more detail below.

Given the CP method outperformed CP-HiL in segmentation efficiency, a quantitative comparison between CP and SSL was made. For this analysis, ten 10x fluorescent images were selected, and 80 cells within these images were randomly chosen for manual segmentation to create a ‘manual mask’. The same cells were then segmented using both CP and SSL methods, generating ‘automated masks’ for comparison. The F1 score was calculated by comparing the ‘manual mask’ against the ‘automated masks’ from each method. The average F1 score for SSL was 0.852 (S.D = ±0.017), surpassing the CP method’s average F1 score of 0.804 (S.D = ±0.08). Representative masks from manual, CP, and SSL segmentation, along with their respective F1 scores, are illustrated in Figure 7. The SSL method demonstrated stable F1 scores across all segmented images, ranging from 0.831 to 0.876. In contrast, the CP method’s F1 scores varied more significantly, from 0.645 to 0.8815, primarily due to misclassified segmentation instances (Figure 6c).

**Figure 6:**
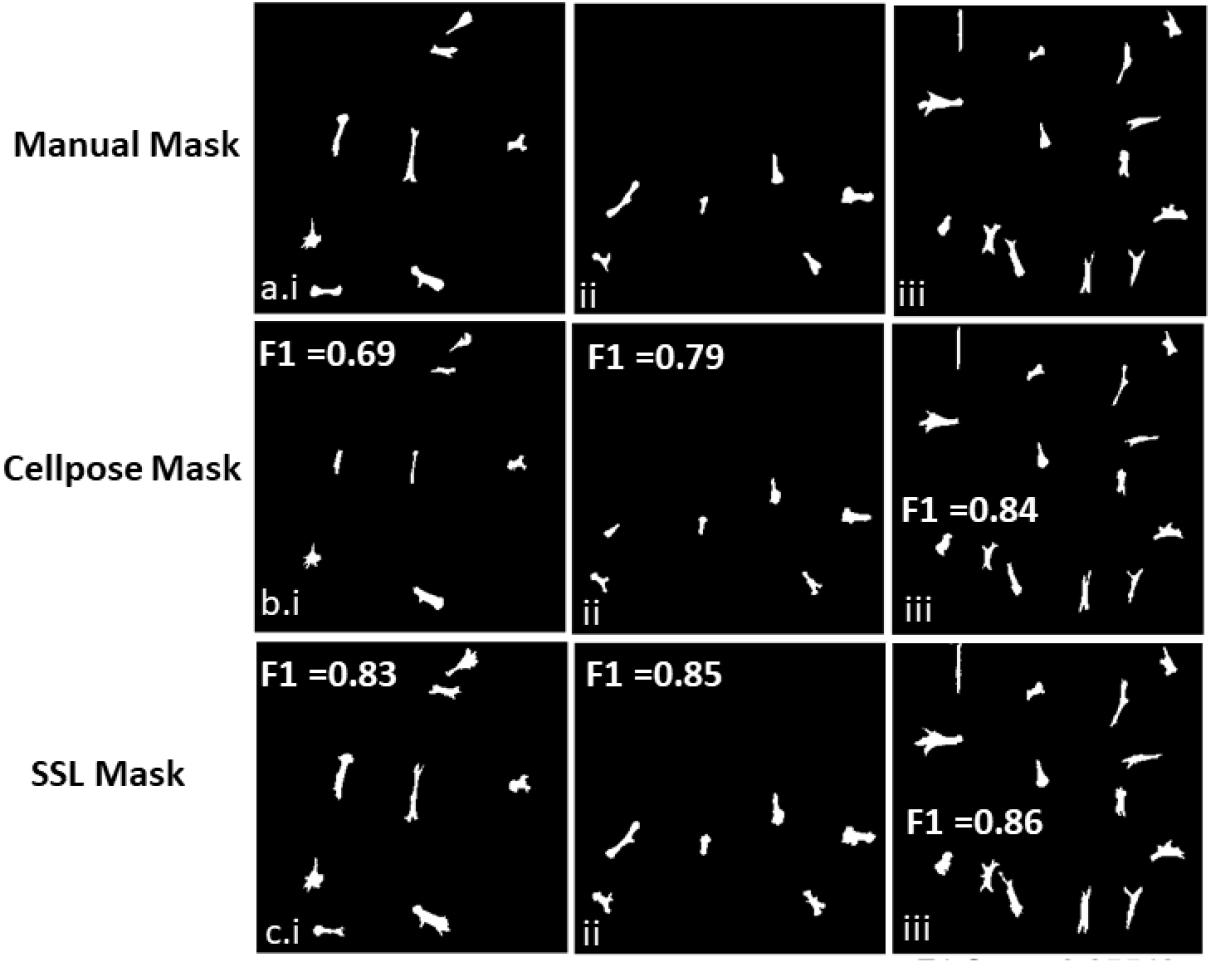
Segmentation masks provided by (a) manual, (b) Cellpose and (c) SSL segmentation.

**Figure 7:**
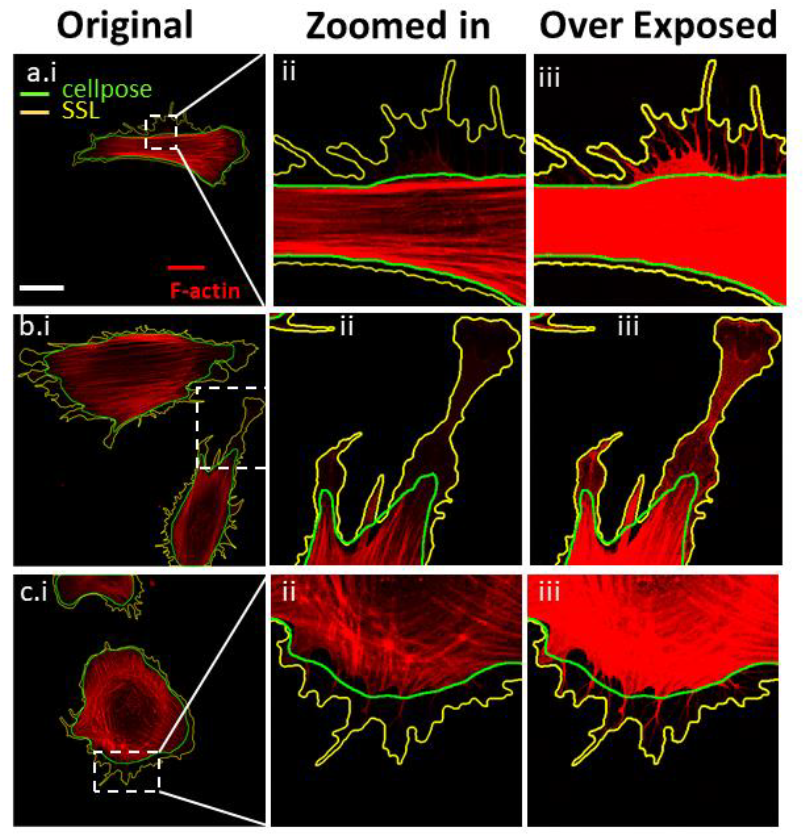
Self-supervised segmentation approach on high-resolution images. Confocal images of Hs27 stained with F-actin for cytoskeleton at 63x objective (a.i, b.i and c.i). Images were zoomed in and over exposed to depict structures of filopodia (a.iii), (c.iii) and lammepodia (b.iii). Scale bar: 10μm.

### Self-supervised learning for enhancing fine-detail cellular structure segmentation

The robustness of SSL in segmenting fine-detail cellular structures represents a significant advancement in high-resolution fluorescent imaging (Figure 7). Our SSL methodologies exhibit a remarkable capacity for accurately delineating intricate cellular components, a task that is essential for detailed biological analysis and understanding. In fact, these structures are not easily seen by eye without contrast enhancements (Figure 7 a.iii, b.iii and c.iii) and were often missed by Cellpose. This precision in segmentation is particularly vital when dealing with complex cellular structures like filopodia (Figure 7a.iii and c.iii) and lamellipodia (Figure 7b.ii), where the distinction between different cellular components can be subtle yet critical.

## Discussion and Conclusion

High throughput *in vitro* imaging is now producing far more imagery - both in terms of number of images and diversity of imaging modalities - than even the most advanced deep learning algorithms can reliably segment. In addition, the increasing requirement of extracting fine cellular details from this imagery means that curated training data sets will need to grow exponentially at a rate that, in our view, is simply unstainable. Machine learning and deep learning pretrained models, when trained on specific cell types or imaging techniques, often do not generalize effectively to other cell types or imaging modalities^22^. Challenges persist due to the inherent variability in cell phenotypes across different tissues and organisms that hampers the creation of universally applicable datasets. Transfer learning can offer a solution to conduct a pre-training model on a similar available dataset and apply it to an unseen target set. Nevertheless, limitations stemming from the pre-training dataset could lead to overfitting, where the model excels with familiar data but underperforms with unexpected phenotypes in the target datasets^30^. This point becomes increasingly more relevant as the popular use of 384 and 1536 well plates in drug discovery applications expands the range of phenotypes. Such wide-ranging imagery datasets can require repeated deep-learning model retraining, making the training procedure increasingly labor and computationally intensive. The innovation of SSL streamlines this process, as each input image generates a specific model for its segmentation purpose, circumventing the pitfalls of over-generalization that often result in reduced accuracy.

An important enhancement to our previous SSL algorithms ^17,18^ is the applicability to single frame data, eliminating the need for live-cell imagery and thus making the code more broadly applicable to include fixed cell imagery. The process of SSL begins with an unsupervised labeling of cell and background pixels by employing optical flow between the original image and an augmented version, thus eliminating the need for curated training data sets and ensuring data relevancy. This method proved effective in identifying intricate organelles and cellular structures regardless of magnification level or the use of external dyes (Figure 3) as well as detecting structures that might be overlooked by the human eye (Figure 7), thereby enhancing the generalizability of machine learning outcomes to high content imaging. The label library generated from the unsupervised step is employed in supervised learning training to classify pixels missed by the optical flow process. The supervised classification in this work leverages three features—entropy, gradient, and intensity—as they provide rich information at the cellular level but is readily expandable. Entropy, a measure of localized texture, is well-known for object edge detection ^31^, and recently applied in a wide range of disciplines like ecology and landscapes^32^, evolutionary biology ^33^ and cancer research ^34–36^. For *in vitro* microscopy, the inherent intracellular variability in organelle and cytoskeletal arrangements make for enhanced texturing versus the background.

Another advantage of this SSL approach is interpretability. The extensive range of data features in deep learning - often requiring extensive tuning of millions of hyperparameters - contributes to its “black box” nature such that scientists can be at a loss to understand the conclusions. This lack of transparency is particularly alarming in biomedical studies ^37^, where understanding the rationale behind a model’s decisions is crucial for advancing discoveries in cellular activity, signaling pathways, or diagnostic procedures. Other pixel classification methods offer many more image features transformed by principal component analysis but their combination can hamper insights ^38,39^. In contrast, our SSL approach utilizes transparent and sufficient known data features focusing on entropy, gradient and intensity without any transformation. Furthermore, the entire SSL application can be operated on a standard laptop computer which avoids any extensive hardware installation and data customization for cloud-based uploading. Our innovative SSL method not only shows improvement in segmentation but also reduces the need of advanced computational skills and risk of data security, thereby lowering the barriers for biological researchers integrating computer vision into their work.

Current studies in cell biology often overlook the meticulous analysis of fine cell structures, instead focusing primarily on basic cellular geometry and fluorescence intensity. This could be either attributed to the limitations of robust and reliable segmenting tools, the imaging tools ^40^ or the data size^41^. The future of cell imaging will need to bridge the gap between high-resolution and high-throughput methodologies. This evolution depends on developing high-throughput cell imaging experiments that are complemented by reliable, straightforward, precise and robust automated cell segmentation techniques. Here we have presented results of our SSL algorithm across various modalities and resolutions (Figure 2, Figure 5c) and experimental treatments (Figure 4) that not only accurately segment cells but also provide detailed structural information (Figure 7) suggesting our code’s ability to meet this need.

Future work will focus on integrating more efficient declumping algorithms for high cell density imagery. Individual cells or cell clump segmentation from the background is a critical step in automated image analysis. Through the application of a watershed algorithm, our current code can effectively delineate borders of MDA-MB-231 (Figures 2c.i) and *S. cervisea* (Figure 2c.ii) when cell concentrations are not overly crowded, but more sophisticated approaches will be needed for higher densities. The segmentation algorithm we have presented here is foundational to all such subsequent analyses, including declumping, morphological measurements, and the extraction of deep learning parameters - all of which are enhanced by first classifying cells from background with high fidelity. Throughout this study, SSL robustly met this goal across various cell types, cell treatments and imaging modalities.

## References

1. Tang, H., Panemangalore, R., Yarde, M., Zhang, L. & Cvijic, M. E. 384-Well Multiplexed Luminex Cytokine Assays for Lead Optimization. SLAS Discov. 21, 548–555 (2016).

2. Knight, S. et al. Enabling 1536-Well High-Throughput Cell-Based Screening through the Application of Novel Centrifugal Plate Washing. SLAS Discov. 22, 732–742 (2017).

3. Bigdelou, P. et al. High-throughput multiplex assays with mouse macrophages on pillar plate platforms. Exp. Cell Res. 396, 112243 (2020).

4. Shariff, A., Kangas, J., Coelho, L. P., Quinn, S. & Murphy, R. F. Automated Image Analysis for High-Content Screening and Analysis. SLAS Discov. 15, 726–734 (2010).

5. Buggenthin, F. et al. An automatic method for robust and fast cell detection in bright field images from high-throughput microscopy. BMC Bioinformatics 14, 297 (2013).

6. Lugagne, J.-B., Lin, H. & Dunlop, M. J. DeLTA: Automated cell segmentation, tracking, and lineage reconstruction using deep learning. PLOS Comput. Biol. 16, e1007673 (2020).

7. Chen, R. Q. et al. Real-time semantic segmentation and anomaly detection of functional images for cell therapy manufacturing. Cytotherapy 25, 1361–1369 (2023).

8. Al-Kofahi, Y., Zaltsman, A., Graves, R., Marshall, W. & Rusu, M. A deep learning-based algorithm for 2-D cell segmentation in microscopy images. BMC Bioinformatics 19, 365 (2018).

9. Caicedo, J. C. et al. Nucleus segmentation across imaging experiments: the 2018 Data Science Bowl. Nat. Methods 16, 1247–1253 (2019).

10. Falk, T. et al. U-Net: deep learning for cell counting, detection, and morphometry. Nat. Methods 16, 67–70 (2019).

11. Long, F. Microscopy cell nuclei segmentation with enhanced U-Net. BMC Bioinformatics 21, 8 (2020).

12. Din, N. U. & Yu, J. Training a deep learning model for single-cell segmentation without manual annotation. Sci. Rep. 11, 23995 (2021).

13. Vuola, A. O., Akram, S. U. & Kannala, J. Mask-RCNN and U-net Ensembled for Nuclei Segmentation. Preprint at http://arxiv.org/abs/1901.10170 (2019).

14. Edlund, C. et al. LIVECell—A large-scale dataset for label-free live cell segmentation. Nat. Methods 18, 1038–1045 (2021).

15. Berg, S. et al. ilastik: interactive machine learning for (bio)image analysis. Nat. Methods 16, 1226–1232 (2019).

16. McQuin, C. et al. CellProfiler 3.0: Next-generation image processing for biology. PLOS Biol. 16, e2005970 (2018).

17. Robitaille, M. C., Byers, J. M., Christodoulides, J. A. & Raphael, M. P. Robust optical flow algorithm for general single cell segmentation. PLOS ONE 17, e0261763 (2022).

18. Robitaille, M. C., Byers, J. M., Christodoulides, J. A. & Raphael, M. P. Self-supervised machine learning for live cell imagery segmentation. Commun. Biol. 5, 1162 (2022).

19. Ji, X., Li, Y., Cheng, J., Yu, Y. & Wang, M. Cell image segmentation based on an improved watershed algorithm. in 2015 8th International Congress on Image and Signal Processing (CISP) 433–437 (IEEE, Shenyang, China, 2015). doi:10.1109/CISP.2015.7407919.

20. Wu, P. et al. Active Contour-Based Cell Segmentation During Freezing and Its Application in Cryopreservation. IEEE Trans. Biomed. Eng. 62, 284–295 (2015).

21. Shen, S. P. et al. Automatic Cell Segmentation by Adaptive Thresholding (ACSAT) for Large-Scale Calcium Imaging Datasets. eneuro 5, ENEURO.0056-18.2018 (2018).

22. Pachitariu, M. & Stringer, C. Cellpose 2.0: how to train your own model. Nat. Methods 19, 1634–1641 (2022).

23. Greenwald, N. F. et al. Whole-cell segmentation of tissue images with human-level performance using large-scale data annotation and deep learning. Nat. Biotechnol. 40, 555–565 (2022).

24. Garvey, C. M. et al. A high-content image-based method for quantitatively studying context-dependent cell population dynamics. Sci. Rep. 6, 29752 (2016).

25. Luengo Hendriks, C. L. et al. [No title found]. Genome Biol. 7, R123 (2006).

26. Zinchenko, V., Hugger, J., Uhlmann, V., Arendt, D. & Kreshuk, A. MorphoFeatures for unsupervised exploration of cell types, tissues, and organs in volume electron microscopy. eLife 12, e80918 (2023).

27. Pieuchot, L. et al. Curvotaxis directs cell migration through cell-scale curvature landscapes. Nat. Commun. 9, 3995 (2018).

28. Robitaille, M. C., Christodoulides, J. A., Calhoun, P. J., Byers, J. M. & Raphael, M. P. Interfacing Live Cells with Surfaces: A Concurrent Control Technique for Quantifying Surface Ligand Activity. ACS Appl. Bio Mater. 4, 7856–7864 (2021).

29. Christodoulides, J. A. et al. Nanostructured substrates for multi-cue investigations of single cells. MRS Commun. 8, 49–58 (2018).

30. Corbe, M., Boncompain, G., Perez, F., Del Nery, E. & Genovesio, A. Transfer learning for versatile and training free high content screening analyses. Sci. Rep. 13, 22599 (2023).

31. El-Sayed, M. A. & Hafeez, T. A.-E. New Edge Detection Technique based on the Shannon Entropy in Gray Level Images. 3, (2011).

32. Cushman, S. Calculation of Configurational Entropy in Complex Landscapes. Entropy 20, 298 (2018).

33. Cushman, S. A. Entropy, Ecology and Evolution: Toward a Unified Philosophy of Biology. Entropy 25, 405 (2023).

34. Lam, V. K., Nguyen, T. C., Chung, B. M., Nehmetallah, G. & Raub, C. B. Quantitative assessment of cancer cell morphology and motility using telecentric digital holographic microscopy and machine learning. Cytometry A 93, 334–345 (2018).

35. Lam, V. K. et al. Machine Learning with Optical Phase Signatures for Phenotypic Profiling of Cell Lines. Cytometry A 95, 757–768 (2019).

36. Vu, T. et al. FunSpace: A functional and spatial analytic approach to cell imaging data using entropy measures. PLOS Comput. Biol. 19, e1011490 (2023).

37. Ching, T. et al. Opportunities and obstacles for deep learning in biology and medicine. J. R. Soc. Interface 15, 20170387 (2018).

38. Elhaik, E. Principal Component Analyses (PCA)-based findings in population genetic studies are highly biased and must be reevaluated. Sci. Rep. 12, 14683 (2022).

39. Chari, T. & Pachter, L. The Specious Art of Single-Cell Genomics. http://biorxiv.org/lookup/doi/10.1101/2021.08.25.457696 (2021) xdoi:10.1101/2021.08.25.457696.

40. Horwath, J. P., Zakharov, D. N., Mégret, R. & Stach, E. A. Understanding important features of deep learning models for segmentation of high-resolution transmission electron microscopy images. Npj Comput. Mater. 6, 108 (2020).

41. Byun, H., Lee, K. & Shim, H. Cell Segmentation in Multi-modality High-Resolution Microscopy Images with Cellpose.

