## Supplementary Note for "A Self-Supervised Learning Approach for High Throughput and High Content Cell Segmentation"

Supplementary Note 1

| Main Text Figure | Cell Type | Optical Modality | Objective Magnification | Objective Numerical Aperture | Camera |
|---|---|---|---|---|---|
| Figure 2a | MDA-MB-231 (human breast adenocarcinoma) | Phase | 10X | 0.3 (air) | Zeiss Axiocam 702 CMOS |
| Figure 2b | MDA-MB-231 (human breast adenocarcinoma) | Bright-field | 40X | oil | |
| Figure 2c.i | MDA-MB-231 (human breast adenocarcinoma) | DIC | 20X | 0.8 (air) | Zeiss Axiocam 702 CMOS |
| Figure 2c.ii | S. cerevisiae | DIC | 63X | 1.4 oil | Zeiss Axiocam 702 CMOS |
| Figure 2d | Hs27 (human foreskin, fibroblast) | IRM | 40X | 1.4 (oil) | Hamamatsu ORCA R2 CCD |
| Figure 2e.i,ii | A549 (human lung adenocarcinoma) | Flourescence (GFP) | 100X | 1.46 oil | Hamamatsu ORCA R2 CCD |
| Figure 2e.iii,iv | MDA-MB-231 (human breast adenocarcinoma) | Flourescence (F-actin and Vinculin) | 60X | 1.25 (oil) | Photometrics CoolSNAP HQ2 CCD |
| Figure 3a | Hs27 (human foreskin, fibroblast) | Flourescence (DAPI, F-actin and Vinculin) | 20X | 0.8 (air) | Zeiss Axiocam 702 CMOS |
| Figure 3b | Hs27 (human foreskin, fibroblast) | Flourescence (F-actin and Vinculin) | 63X | | |
| Figure 4 | Hs27 (human foreskin, fibroblast) | Fluorescence (Zeiss3) (Vinculin) | 40X | 1.4 (oil) | Hamamatsu ORCA R2 CCD |
| Figure 5a.i-ii, b.i-ii, c.i-iii | Hs27 (human foreskin, fibroblast) | Fluorescence (F-actin) | 10X | 0.3 (air) | Zeiss Axiocam 702 CMOS |
| Figure 5a.iv-vi, b.iv-vi, c.iv-vi | Hs27 (human foreskin, fibroblast) | Flourescence (F-actin) | 63X | | |

Supplementary Note 2: Self-supervised learning segmentation approach before and after applying declumping step

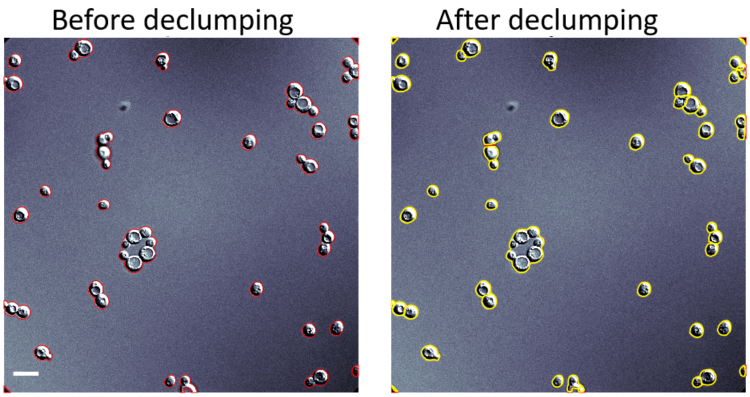


Figure: Self-supervised learning segmentation approach on S. cerevisiae image taken by DIC (63X objective) with and without declumping step. Scale bar: 5µm

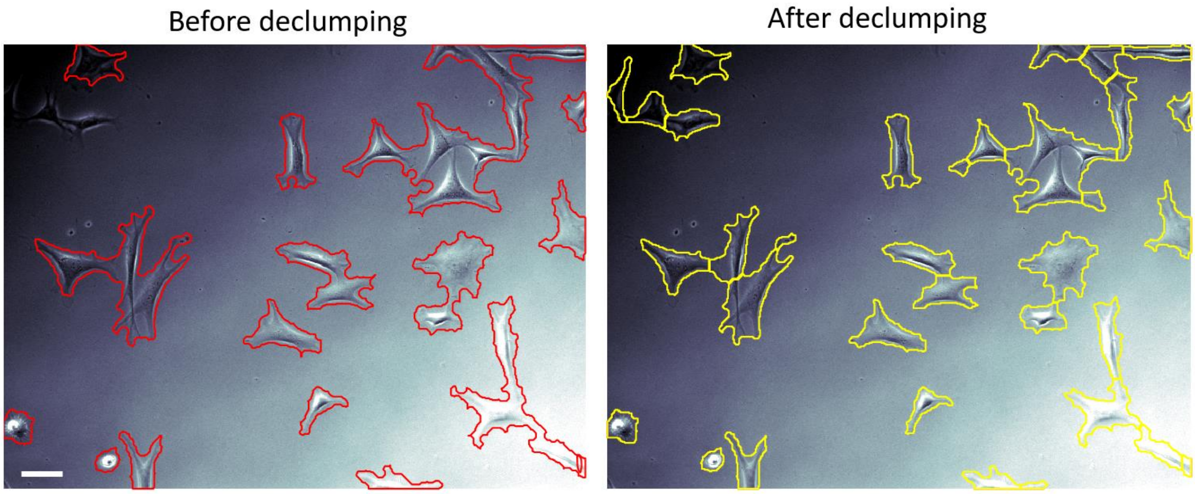


Figure: Self-supervised learning segmentation approach on Hs27 image taken by phase microscopy (10X objective) with and without declumping step. Scale bar: 50µm
